# LRRK2-phosphorylated Rab10 sequesters Myosin Va with RILPL2 during ciliogenesis blockade

**DOI:** 10.1101/2020.04.28.065664

**Authors:** Izumi Yanatori, Herschel S. Dhekne, Edmundo G. Vides, Yuriko Sobu, Federico Diez, Suzanne R. Pfeffer

## Abstract

Activating mutations in LRRK2 kinase cause Parkinson’s disease. Pathogenic LRRK2 phosphorylates a subset of Rab GTPases and blocks ciliogenesis. Thus, defining novel phospho-Rab interacting partners is critical to our understanding of the molecular basis of LRRK2 pathogenesis. RILPL2 binds with strong preference to LRRK2-phosphorylated Rab8A and Rab10. RILPL2 is a binding partner of the motor protein and Rab effector, Myosin Va. We show here that the globular tail domain of Myosin Va also contains a high affinity binding site for LRRK2-phosphorylated Rab10, and certain tissue-specific Myosin Va isoforms strongly prefer to bind phosphorylated Rab10. In the presence of pathogenic LRRK2, RILPL2 relocalizes to the peri-centriolar region in a phosphoRab10- and Myosin Va-dependent manner. In the absence of phosphoRab10, expression of RILPL2 or depletion of Myosin Va increase centriolar RILPL2 levels, and either condition is sufficient to block ciliogenesis in RPE cells. These experiments show that LRRK2 generated phosphoRab10 dramatically redistributes Myosin Va-RILPL2 complexes to the mother centriole, which may sequester Myosin Va and RILPL2 in a manner that blocks their normal roles in ciliogenesis.

## Introduction

Activating mutations in the LRRK2 kinase cause Parkinson’s disease (Alessi & Sammler, 2018). Recent work has shown that pathogenic LRRK2 phosphorylates a subset of Rab GTPases, including Rabs 8,10,12 and 35 (Steger *et al*, 2017, 2016) that are master regulators of membrane trafficking events (Pfeffer, 2017). A significant consequence of LRRK2 Rab phosphorylation is a decrease in primary cilia in a variety of cell types, including in mouse brain (Steger *et al*, 2017; Dhekne *et al*, 2018; Lara Ordónez *et al*, 2019). Phosphorylation of Rab proteins blocks their abilities to interact with key regulators including Rab GDI and the Rabin8 guanine nucleotide exchange factor, as well as multiple, cognate Rab effector proteins (Steger *et al*, 2016). This loss of effector binding alone would be sufficient to interfere with Rab protein physiological functions. Importantly, however, phosphorylated Rab proteins also show enhanced binding to novel effectors, and understanding the roles of these nascent interactions is critical to our understanding of the molecular basis of Parkinson’s disease pathogenesis.

RILP is a Rab7 effector and so-called, “cargo adaptor” that links the microtubule-based motor proteins, dynein/dynactin to late endosomes (Cantalupo *et al*, 2001; Jordens *et al*, 2001). RILPL1 and RILPL2 are two RILP-related proteins that bind much more tightly to phosphorylated Rab8 and Rab10 proteins than to their non-phosphorylated states (Steger *et al*, 2017), but little is known about their cellular roles. Both RILPL1 and RILPL2 were reported to be ciliary proteins involved in regulating ciliary content (Schaub & Stearns, 2013). We showed previously that RILPL1 is relocalized quantitatively to the mother centriole by (phosphorylated) pRab10 protein, and both pRab10 and RILPL1 (but not Rab8A) are essential for the ability of pathogenic LRRK2 to block cilia formation (Dhekne *et al*, 2018). In this study, we have investigated the possible role of RILPL2 in contributing to a LRRK2-triggered cilia blockade.

RILPL2 is a Myosin Va (MyoVa)-interacting protein first reported to be important for cell shape control and neuronal morphogenesis (Lisé *et al*, 2009). RILPL2 interacts with the C-terminal, globular tail domain of MyoVa via its N-terminal “RH1” domain (Lisé *et al*, 2009) and binds pRab8 and pRab10 (Steger *et al*, 2017) via its C-terminal RH2 domain (129-165; cf. Waschbüsch *et al*, 2020). Recently, RILPL2 has also been reported to regulate the motor activity of MyoVa (Cao *et al*, 2019). By binding a myosin motor at one end, and a membrane-anchored Rab at the other end, RILPL2 is an actin-based, motor-adaptor protein.

We have been studying the effect of Rab phosphorylation on ciliogenesis and report here our progress in understanding the consequences of pRab10-RILPL2 complex formation. To our surprise we find that the MyoVa globular tail domain also contains a binding site for LRRK2-phosphorylated Rab10, and LRRK2 kinase activity and pRab10 drive MyoVa and RILPL2 to the mother centriole during ciliogenesis blockade, perhaps entrapping MyoVa at that location.

## Results

### RILPL2 re-localizes to the mother centriole in serum starved cells

RILPL2 associates with primary cilia in IMCD3 cells (Schaub & Stearns, 2013). We determined the localization of RILPL2 in primary astrocytes that were isolated using antibody immuno-panning from brains of newborn rats (Foo *et al*, 2011). Most of these astrocytes contain Arl13b^+^ primary cilia, and endogenous RILPL2 was detected at low levels throughout the cytoplasm and also at the base of these cilia (Fig. 1A). Cultured cell lines including hTERT-RPE cells yielded only weak endogenous RILPL2 staining, however upon exogenous expression, HA-RILPL2 displayed a diffuse, cytoplasmic localization (see below). After 2 hours of serum starvation to induce primary cilia formation, a small amount of HA-RILPL2 was detected adjacent to the mother centriole, identified by distal appendage protein CEP164, centriolar cap protein CP110 or the ciliary vesicle marker, EHD1 (Fig. 1B); partial RILPL2 re-localization upon serum withdrawal was observed in >50% of the transfected cells (Fig. 1C).

**Figure 1.**
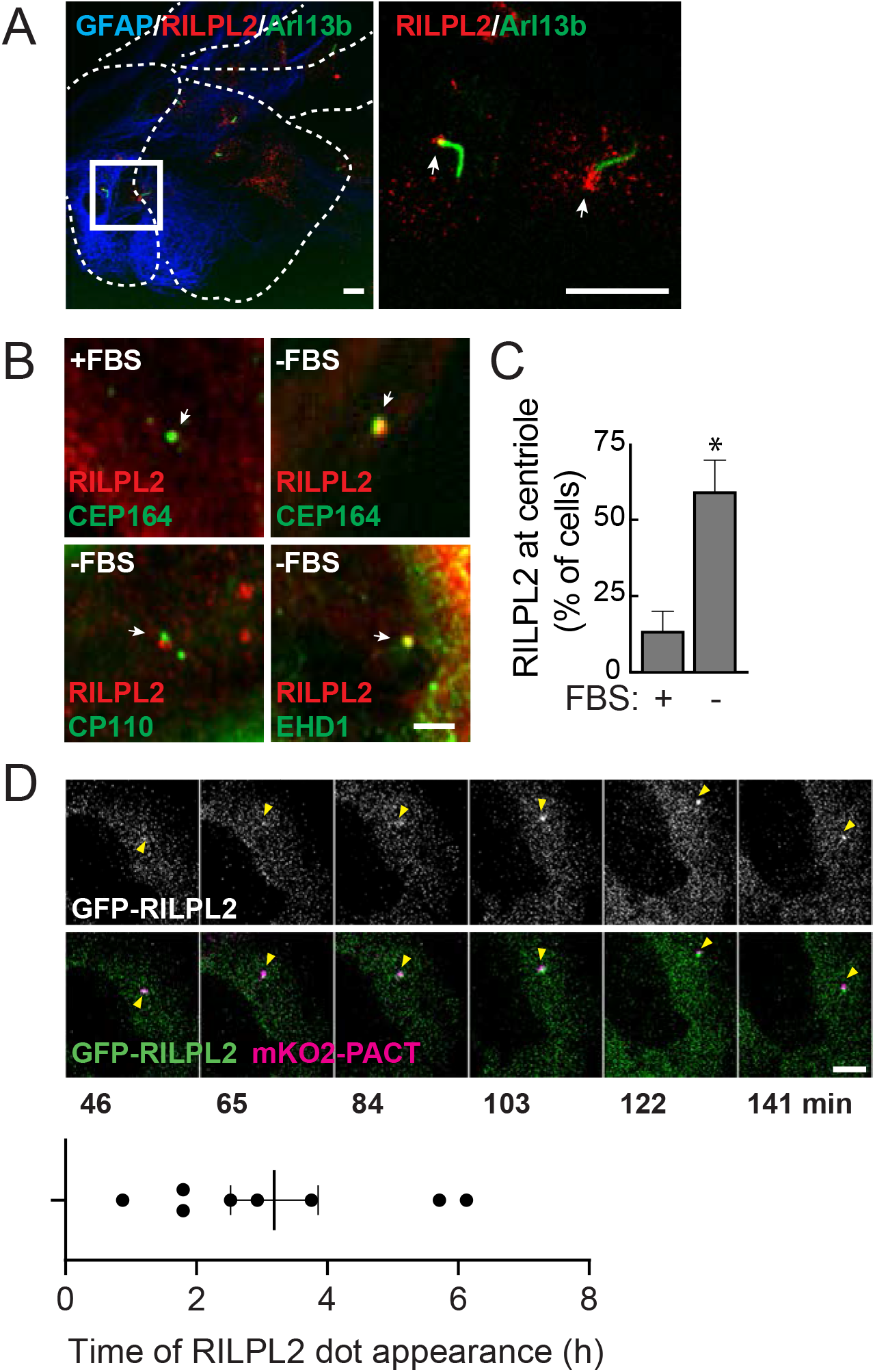
RILPL2 re-localizes to the mother centriole in serum starved cells. (A) Primary rat astrocytes were fixed with 3% PFA and stained for cilia using mouse anti-Arl13b (green) and rabbit anti-RILPL2 (red). Box indicates the cropped region of interest enlarged in the image shown at right. Bar, 10 μm. (B) hTERT-RPE cells were transfected with HA-RILPL2 at 80% confluency and 24 h later either medium changed or serum starved for 2 h. Cells were fixed with −20°C methanol for 2 min and stained with rabbit or mouse anti-HA to detect RILPL2 (red) and mouse anti-CEP164, rabbit anti-CP110 or rabbit anti-EHD1 (green). Presence or absence of FBS is indicated. Arrows indicate the location of the centriolar region. Bar, 2.5 μm. (C) Quantitation of percent of cells that show centriolar localized RILPL2 ± 2h serum starvation. Significance was determined by the t-test; *, p = 0.03. Error bars represent standard error of the mean from two independent experiments with >50 cells per condition. (D) hTERT-RPE cells expressing mKO2-PACT (magenta) were transfected with GFP-RILPL2 (green). Cells were serum starved and 15 min later imaged by capturing Z-stacks every 6 min for the next 8 hours. Yellow arrowheads indicate the location of the centrosome as marked by mKO2-PACT. Upper panel shows GFP-RILPL2 in grayscale. Bottom panel shows the times at which GFP-RILPL2 appeared at the mKO2-PACT structure. Error bar represents standard error of the mean. Scale bar, 5 μm.

Upon live imaging of hTERT-RPE cells transfected with GFP-RILPL2 and mKO2 tagged with the centrosomal targeting sequence of Pericentrin/AKAP450 (PACT) to provide a centrosomal marker (Gillingham & Munro, 2000), RILPL2 arrived at the mKO2-PACT^+^ puncta after ∼3 hours of serum starvation, and remained associated with the centrosome for at least the next 5 hours (Fig. 1D).

### MyoVa regulates centriolar RILPL2 localization

As mentioned earlier, RILPL2 binds MyoVa. Most MyoVa is distributed diffusely throughout the cytoplasm (see below) but a small amount of total cellular MyoVa associates with primary cilia (Kohli *et al*, 2017) and regulates ciliogenesis (Assis *et al*, 2017). Wu *et al* (2018) reported that MyoVa mediates transport of preciliary vesicles from the peri-centrosomal region to the distal appendages of the mother centriole. To explore the possible contribution of MyoVa to centriolar RILPL2 localization, we depleted MyoVa using siRNA and then monitored the localization of exogenously expressed HA-RILPL2 (Figs. 2A, B). To our surprise, RILPL2 localized to the mother centriole in >50% of MyoVa-depleted cells, even under non-ciliating conditions, compared with ∼15% of control siRNA-transfected cells (Figs. 2C, D). These data indicate that MyoVa transports RILPL2 away from the centriolar region under non-ciliating conditions.

**Figure 2.**
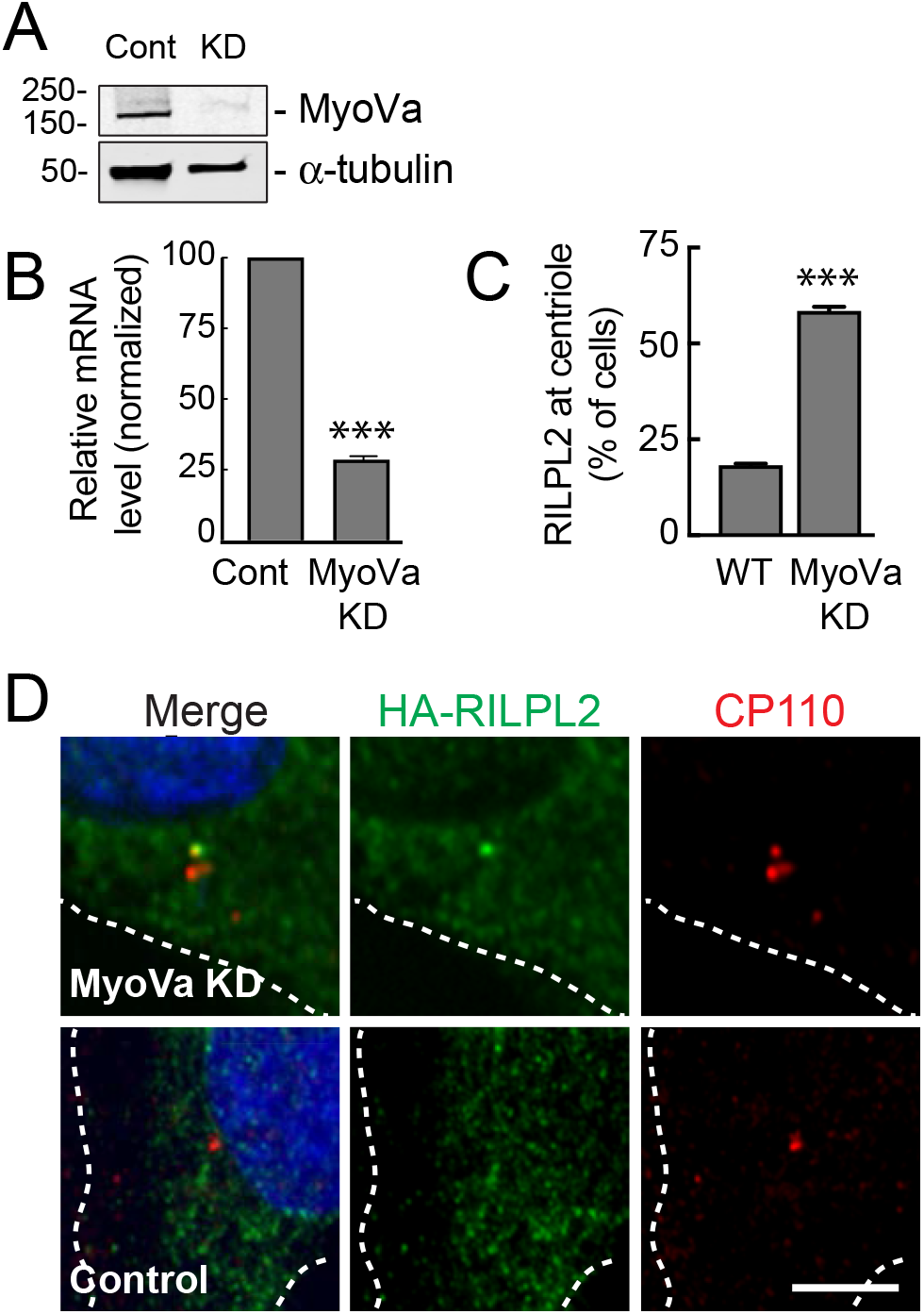
MyoVa regulates centriolar localization of RILPL2. A 6-well plate of hTERT-RPE cells was transfected with siRNA specific for MyoVa (MyoVa-KD) or control siRNA (cont) and 48 h later assayed for mRNA levels by qPCR and 72 h later for protein levels by MyoVa western blot. (A) Immunoblot of hTERT-RPE cells transfected as indicated and 72 h later lysed in NP40 lysis buffer. SDS-PAGE of 60μg protein lysate and immunoblot performed using rabbit anti-MyoVa and mouse anti-tubulin antibodies. Numbers adjacent to the gels (in this and all subsequent figures) indicate the molecular mass markers in kDa. (B) Graph indicates relative mRNA level by qPCR analysis of MyoVa normalized to GAPDH in control (cont) and MyoVa (MyoVa KD) si-RNA treated cells. Significance was determined by the t-test; ***, p = 0.0004. (C,D) SiRNA transfected cells were transfected with HA-RILPL2 for 24 h. Cells were methanol fixed and stained with mouse anti-HA (green) and rabbit anti-CP110 (red). Dotted lines indicate the cell boundaries. Bar, 5 μm. (C) Quantitation of cells that have RILPL2 localized at the mother centriole in control (WT) and MyoVa (MyoVa KD) si-RNA-treated conditions. Error bars indicate the standard error of the mean from two independent experiments. Significance was determined by the t-test; ***, p = 0.0005 from 2 independent experiments with >25 cells counted.

Note that both N-terminally HA- and GFP-tagged RILPL2 were detected at the centriole upon serum starvation (Figs. 1B, D) or MyoVa depletion (Fig. 2D), however less C-terminally tagged RILPL2-GFP localized to the centriolar region in parallel experiments, suggesting a role for the RILPL2 C-terminus in centriolar localization, as reported previously (Schaub and Stearns, 2013).

### MyoVa colocalizes with pRab10 independent of RILPL2

Like RILPL2, the related RILPL1 also binds preferentially to phosphorylated Rab8 and Rab10 (Steger *et al*, 2017; Waschbüsch *et al*, 2020). We showed previously that RILPL1 enhances the pericentriolar clustering of pRab10-containing membrane compartments (Dhekne *et al*, 2018). We therefore explored the relationship between RILPL2 expression and pRab10 distribution. In cells expressing R1441G LRRK2 to generate pRab10, RILPL2 was cytosolic (Red, Fig. 3A) and a fraction (∼20%) was concentrated in the peri-centriolar region with pRab10 (Green, Figs. 3A, B). pRab10 was significantly more concentrated in the centriolar region than RILPL2 under these conditions.

**Figure 3.**
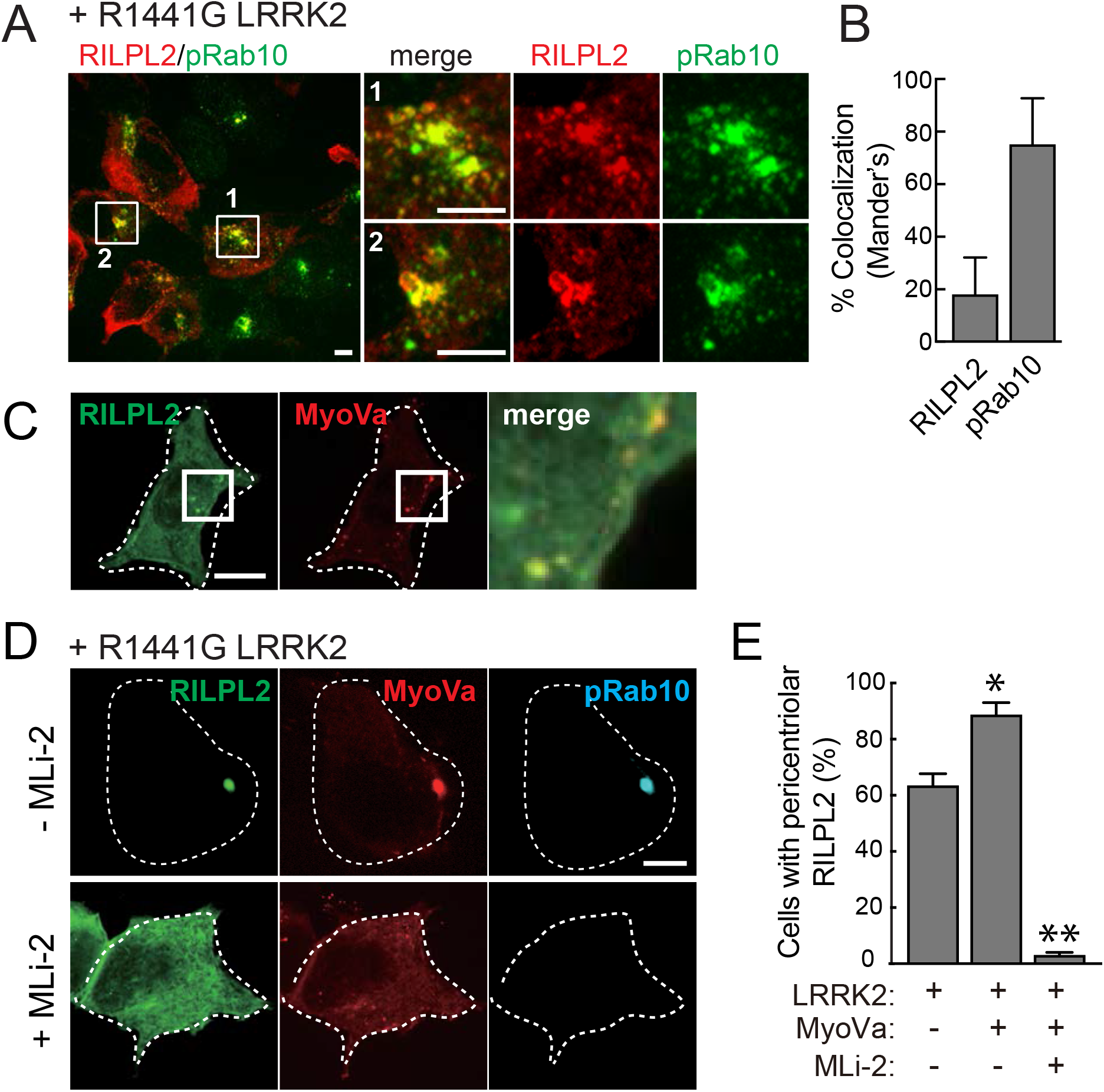
Pathogenic LRRK2 drives RILPL2 and MyoVa to cluster with pRab10. (A) HEK293T cells were co-transfected with myc-LRRK2-R1441G and HA-RILPL2. After 18h, cells were fixed and stained with rabbit anti-pRab10 (green) and mouse anti-HA (red). Bar, 10 μm. Boxes indicate the regions that are cropped and expanded at right. Regions 1 and 2 are as indicated. Bars, 5 μm. (B) Quantitation of colocalization using Mander’s cooccurrence method. RILPL2 indicates the fraction of total RILPL2 that colocalizes with pRab10. pRab10 indicates the fraction of total pRab10 that colocalizes with RILPL2. Error bars indicate standard error of the mean for >20 cells. (C) HEK293T cells were cotransfected with RILPL2-GFP, MyoVa-mCherry. Cells were fixed with 3% PFA. RILPL2 (green) and MyoVa (red) are indicated. Bar, 10 μm. (D) HEK293T cells were co-transfected with RILPL2-GFP, MyoVa-mCherry ± Myc-LRRK2-R1441G. After 18h, LRRK2 transfected cells were treated with 200nM MLi-2 or DMSO for 2h. Cells were fixed and stained for pRab10 (cyan). MLi-2 treatment is indicated. Dotted outlines indicate cell boundaries. RILPL2 (green) and MyoVa (red) are indicated. Bar, 10 μm. (E) Quantitation of cells with pericentriolar RILPL2 in the presence of transfected LRRK2, co-transfected with MyoVa or treated with MLi-2 as indicated. Error bars indicate standard error of the mean from two independent experiments with >50 cells per condition. Significance was determined by the t-test; *, p = 0.027; **, p = 0.0026.

Upon co-transfection in control cells, RILPL2 and MyoVa showed minimal colocalization that was limited to a few distinct puncta, and most RILPL2 protein was cytosolic (Fig. 3C). Introduction of pathogenic R1441G LRRK2 caused essentially all of the MyoVa and RILPL2 proteins to concentrate in the peri-centriolar region and to colocalize in more than 80% of the cells (Fig. 3D, top row). Importantly, addition of the MLi-2 LRRK2 inhibitor for two hours completely reversed the phenotype; under these conditions, pRab10 staining was lost and both RILPL2 and MyoVa proteins reverted to a diffuse localization in 90% of cells (Figs. 3D, E). Thus, in the absence of LRRK2, MyoVa appears to disperse RILPL2 but in the presence of pathogenic LRRK2 which generates pericentriolar pRab10, RILPL2 and MyoVa strongly relocalize to the mother centriole.

### Rab10 drives LRRK2-dependent relocalization of MyoVa and RILPL2

MyoVa binds both Rab10 and RILPL2 (Lisé *et al*, 2009; Roland et al, 2009). Without exogenous RILPL2 co-expression (unlike Fig. 3C), R1441G LRRK2 activity led to the relocalization of ∼40% MyoVa to the pericentriolar region where it co-localized with pRab10 (Fig. 4A). Under these conditions, pRab10 was highly concentrated at the centriole and >80% of pRab10 colocalized with MyoVa (Fig. 4B). LRRK2 phosphorylates multiple Rab GTPases in addition to Rab10 including Rabs 8, 12, 35 and 43 (Steger *et al*, 2017). Nevertheless, Rab10 was absolutely required for the LRRK2-triggered relocalization of MyoVa or RILPL2 tested independently, as neither protein re-localized to the centriole in cells depleted of Rab10 or treated with LRRK2 inhibitor, MLi-2 (Figs. 4C-E).

**Figure 4.**
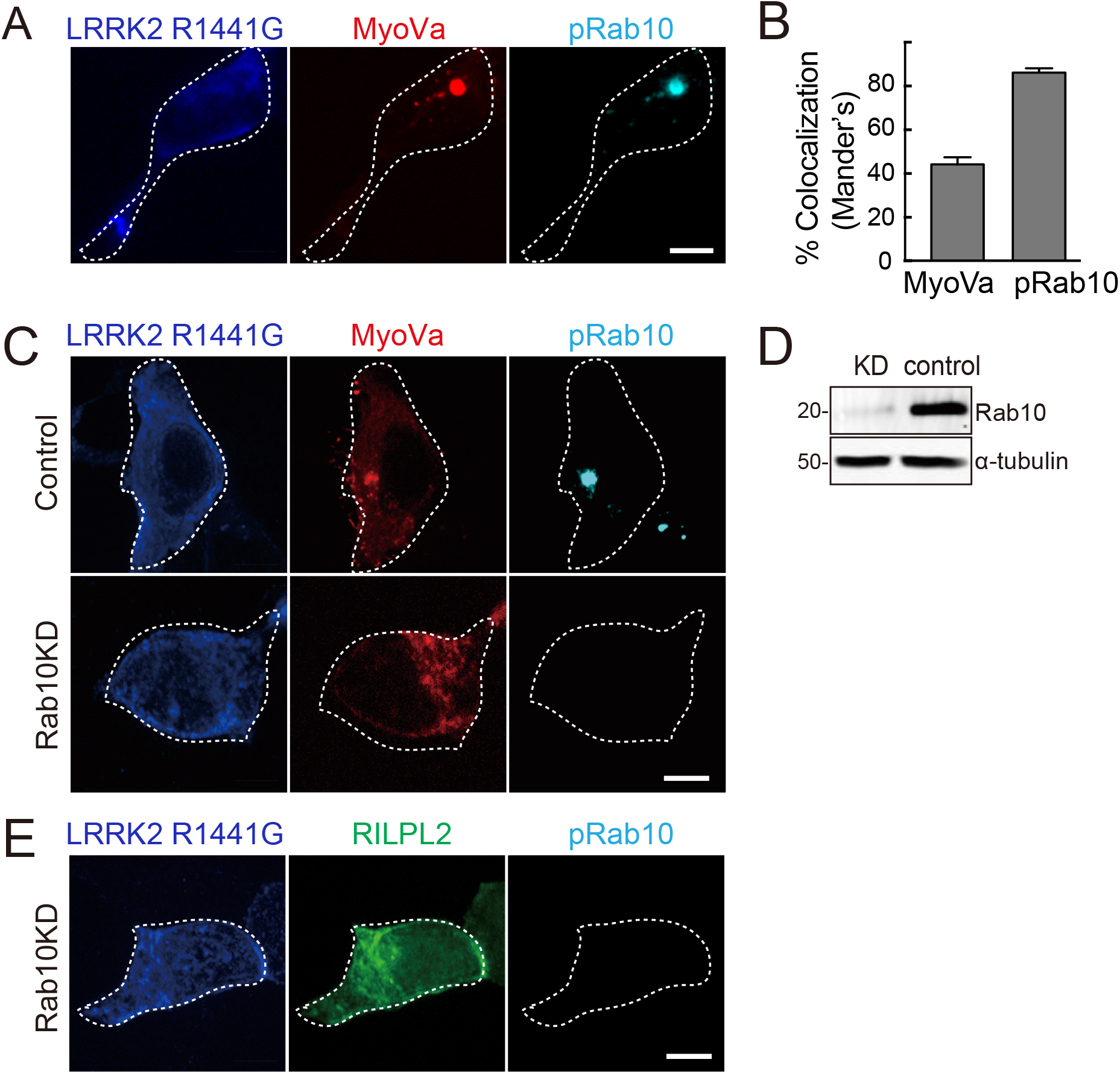
Pathogenic LRRK2 drives MyoVa colocalization with pRab10 via Rab10. (A) HEK293T cells were co-transfected with Myc-LRRK2-R1441G and MyoVa-mCherry. After 18h, cells were fixed and stained with mouse anti-Myc (blue) and rabbit anti-pRab10 (cyan). MyoVa-mCherry (red) is indicated. (B) Quantitation of colocalization by Mander’s method between MyoVa and pRab10. The fraction of MyoVa that colocalizes with pRab10 and vice-versa are quantified. Error bars indicate standard error of the mean of colocalization coefficient for >20 cells. (C) HEK293T cells depleted of Rab10 (KD) or control cells were cotransfected with Myc-LRRK2 R1441G and MyoVa-mCherry. After 18h, cells were stained with mouse anti-Myc (blue) and rabbit anti-pRab10 (cyan). MyoVa-mCherry (red) is indicated. (D) HEK293T cells were infected with lentiviral shRNA targeting Rab10 to create a knock-down (KD) or scrambled control and selected using puromycin. Cells were lysed in NP40 lysis buffer and 60 μg lysate was immunoblotted for Rab10 with tubulin as loading control. (E) HEK293T cells that are knocked-down for Rab10 were co-transfected with myc-LRRK2 R1441G and RILPL2-GFP. After 18 h, cells were stained with mouse anti-myc (blue) and rabbit anti pRab10 (cyan). RILPL2-GFP (green) is indicated. Rab10 knock down (Rab10KD) is indicated. Outlines indicate the boundaries of the cells being shown. Bars, 10 μm.

### pRab10 binding to the MyoVa globular tail domain

Figure 5A diagrams the domain structure of Myosin V proteins. The N-terminal motor domain is followed by a series of IQ motifs and a set of variably spliced exons. At the C-terminus, the MyoVa globular tail domain (GTD) can interact with Rab3A (Wollert *et al*, 2011), Rab11 (Roland *et al*, 2009; Lindsay *et al*, 2013; Pylypenko *et al*, 2013, 2016) and with Rab27A via the Slac2/Melanophilin adaptor (Fukuda *et al*, 2002; Hume *et al*, 2002; Wei *et al*, 2013) in different cell types. Importantly, Exon D that is conserved in all Myosin Vs contains a binding site for Rab10 (Roland *et al*, 2009). Exon D-containing forms of MyoVa are expressed broadly, but not in brain or endocrine/neuroendocrine cells (Seperack et al, 1995; Roland *et al*, 2009). We therefore created a MyoVa truncation construct that deletes Exon D (MyoVa-ΔD lacking amino acids 1320-1345) or enables expression of the GTD alone (residues 1421 to 1880; Fig. 5A).

**Figure 5.**
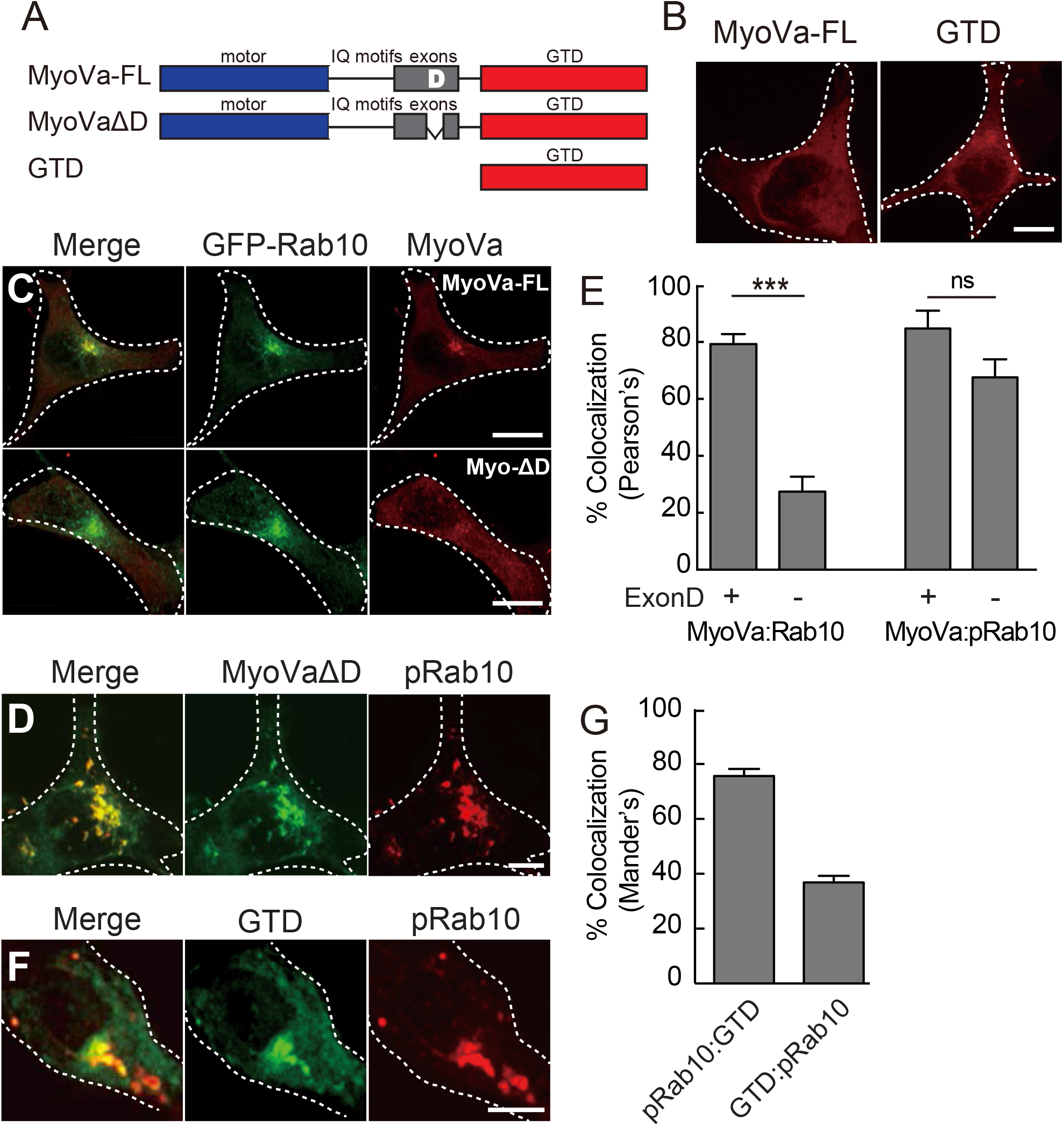
MyoVa relies on exon D and its globular tail domain for Rab10 colocalization. (A) Diagram of MyoVa domain structure. MyoVa has a motor domain (blue), isoleucine-glutamine (IQ) motif, five exons (A-E, gray) and a globular tail domain (GTD, red). MyoVa ΔD lacks exon D; GTD indicates a construct that contains only the MyoVa C-terminus. (B) Full length MyoVa-mCherry (MyoVa-FL) or GTD-mCherry were transfected into HEK293T cells plated on collagen coated coverslips and imaged after 24h. Bars, 10 μm. (C) GFP-Rab10 was co-transfected with full length MyoVa-mCherry (top row) or ExonD deleted MyoVa-mCherry (bottom row) in HEK293T cells and fixed after 24h. Bar, 10 μm. (D) Myc-LRRK2 R1441G was co-transfected with Exon D deleted-MyoVa-mCherry in HEK293T cells and fixed after 24h. Cells were stained for pRab10 (red) while MyoVa mCherry is pseudo-colored (green). Bar, 10 μm. (E, left) Quantitation of colocalization by Pearson’s method between GFP-Rab10 and MyoVa-mCherry ± Exon D (as indicated). (E, Right) Quantitation of colocalization by Pearson’s method between pRab10 and MyoVa-mCherry ± Exon D (as indicated). (F) Myc-LRRK2 R1441G was co-transfected with GTD-mCherry in HEK293T cells and fixed after 24h. Cells were stained for pRab10 (red) while GTD-mCherry is pseudocolored (green). Bar, 10 μm. (G) Quantitation of colocalization by Mander’s co-occurrence method where fraction pRab10 that colocalizes with GTD is pRab10:GTD and fraction of GTD that colocalizes with pRab10 is GTD:pRab10. Scale bars, 10 μm. Error bars indicate standard error of the mean from three independent experiments with >50 cells per condition. Significance was determined by the t-test; ***, p = 0.0001; ns = not significant with p = 0.0673.

In HEK-293T cells, mCherry-tagged versions of MyoVa-FL (full length MyoVa) and GTD were mostly cytosolic (Fig. 5B). When MyoVa-FL was co-expressed with GFP-Rab10, more of it re-localized to perinuclear, Rab10 containing vesicles (Fig. 5C), consistent with its established binding capacity. In contrast, Exon D-deleted MyoVa (MyoVa-ΔD) showed much less colocalization with GFP-Rab10 (80 versus ∼30%, Figs. 5C, E). Surprisingly, expression of R1441G pathogenic LRRK2 relocalized both MyoVa-FL (Fig. 4C) and the MyoVa-ΔD to perinuclear membrane structures positive for pRab10 (Fig. 5D). Quantitation of colocalization revealed that MyoVa colocalized with pRab10 to almost the same extent, with or without Exon D (Fig. 5E). Importantly, even the normally diffuse GTD (Fig. 5B) redistributed to pericentriolar, pRab10^+^ membranes (Fig. 5F): 35% of total MyoVa GTD colocalized with pRab10, while ∼80% of pRab10 co-localized with GTD (Fig. 5G). These values match closely the extent of RILPL2 relocalization seen in earlier experiments (Fig. 3).

The ability of pRab10 to relocalize both MyoVa-ΔD and the GTD strongly suggested that the MyoVa GTD contains a pRab10-specific binding site. This was tested in co-immunoprecipitation experiments in cells expressing R1441G LRRK2. As shown in Fig. 6, full length MyoVa-mCherry bound both Rab10 and pRab10; binding of both forms decreased in the presence of MLi-2 LRRK2 inhibitor, suggesting that pRab10 was a significant contributor to MyoVa binding. Similarly, MyoVa-ΔD lacking the D exon-Rab10 binding site still bound roughly the same amount of pRab10 and less total Rab10 protein (Figs. 6A,B). These experiments strongly suggest that MyoVa-ΔD contains an additional binding site with preference for pRab10. Indeed, similar analysis of binding to the GTD domain in cell extracts (Fig. 6C) confirmed co-immunoprecipitation of Rab10 and pRab10 binding to the GTD, both of which were completely lost upon LRRK2 inhibition.

**Figure 6.**
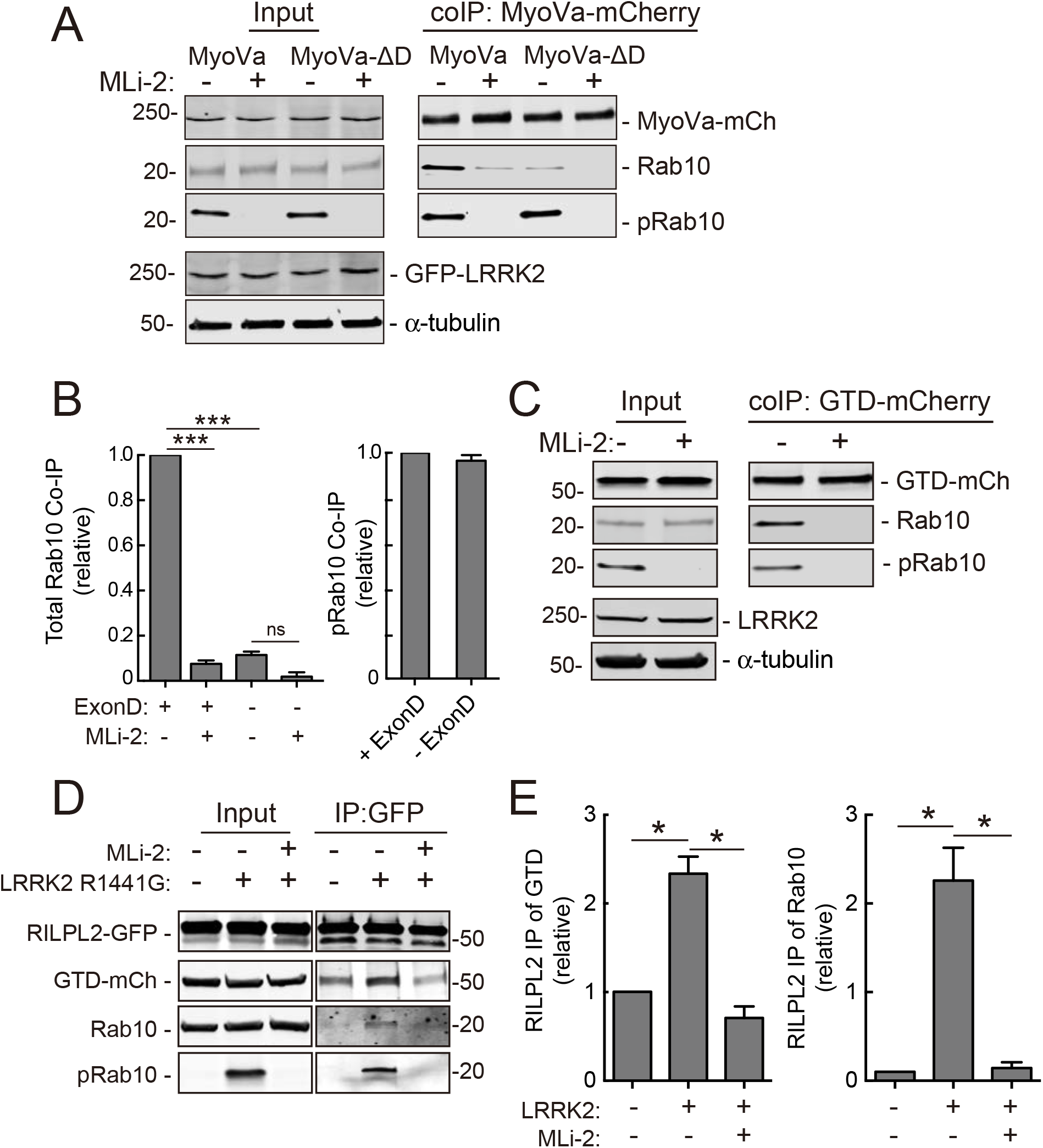
pRab10-specific interaction with the globular tail domain IP. (A) MyoVa- or MyoVa ΔD-mCherry and GFP-LRRK2-R1441G were co-transfected in HEK293T cells. 24h post transfection, cells were incubated with 200 nM MLi-2 for 4h. Cells were lysed in lysis buffer and 400μg extract immunoprecipitated with anti-RFP antibodies on protein G beads. Samples (50% of IP and 15% of input for all IPs in all panels) were immunoblotted for anti-GFP, anti-RFP, anti-tubulin, anti-Rab10 and anti-pRab10 antibodies. (B) Quantitation of the relative amount of Rab10 (left) or pRab10 (right) coimmunoprecipitated by MyoVa and mutant in (A). Error bars indicate standard error of the mean from two gels per co-IP. Significance was determined by t-test; ***, p = 0.0003; ns = not significant with p = 0.295. (C) MyoVa GTD-mCherry and GFP-LRRK2 R1441G were cotransfected into HEK293T cells. After 24h, cells were incubated with 200 nM MLi-2 for 4 h. Cells were lysed in lysis buffer and immunoprecipitated with anti-RFP antibodies on protein G beads. Samples were immunoblotted for anti-GFP, anti-RFP, anti-Rab10 and anti-pRab10 antibodies. (D) RILPL2-GFP, MyoVa GTD-mCherry and Myc-LRRK2 R1441G were cotransfected into HEK293T cells. 24h post-transfection, cells were incubated with 200 nM MLi-2 for 4 h. Cells were lysed in lysis buffer and immunoprecipitated with GFP binding protein-Sepharose. Samples were immunoblotted for anti-GFP, anti-RFP, anti-Rab10 and anti-pRab10 antibodies. (E) Quantitation of the relative amount of the total MyoVa GTD (left) and Rab10 (right) co-immunoprecipitated by RILPL2-GFP (D). Significance was determined by the t-test; *, p = 0.0207 (left) and p = 0.03 (right).

In experiments in which we monitored binding to RILPL2-GFP, MyoVa-GTD bound with or without LRRK2 activity, but we detected about twice as much MyoVa-GTD and Rab10 precipitating with RILPL2 in cells expressing R1441G LRRK2 (Figs. 6D,E). These data are consistent with pRab10 binding both RILPL2 and MyoVa, perhaps forming a ternary complex. Since RILPL2 can dimerize (Wei *et al*, 2013; Waschbüsch *et al*, 2020), it is possible that each half of the dimer engages either MyoVa or pRab10 or both. Alternatively, pRab10 may enhance the ability of MyoVa to bind RILPL2.

### Purified MyoVa globular tail domain binds phosphorylated Rab10 directly

To demonstrate more directly the presence of a pRab10-specific binding site in the MyoVa GTD, we used microscale thermophoresis (MST) to monitor binding of purified proteins. Bacterially expressed, purified Rab10 was phosphorylated in vitro using Mst3, a promiscuous kinase used previously to phosphorylate purified Rab8 and Rab10 proteins (Berndsen *et al*, 2019; Waschbüsch *et al*, 2020). In vitro phosphorylation of Rab10 proceeded linearly for ∼100 minutes before reaching a plateau in reactions containing 2mM ATP and a ratio of 1:3 kinase:substrate at 27°C, as detected using a phosphoRab10-specific antibody (Fig. 7A). As expected, the reaction was ATP dependent (Fig. 7B), and phosphorylation was efficient: ∼90% of Rab10 was phosphorylated in reactions containing Mst3 kinase as determined by PhosTag gel electrophoresis that enables resolution of phosphorylated and nonphosphorylated Rab10 proteins (Fig. 7C).

**Figure 7.**
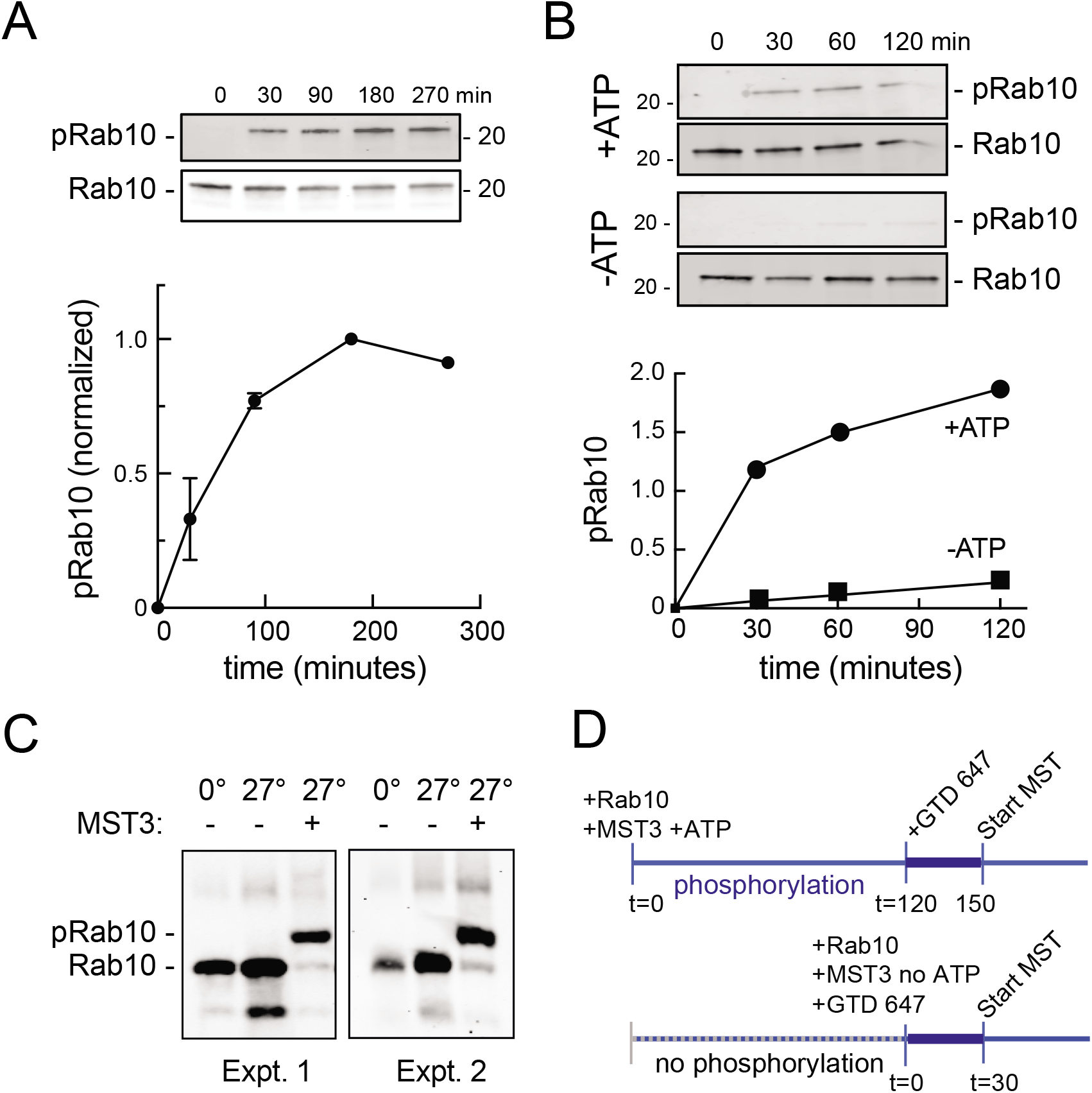
In vitro phosphorylation of Rab10 by Mst3 kinase. (A) Immunoblot of reactions containing purified Rab10 Q63L and Mst3, incubated together at 27°C with 2 mM ATP. Reactions were stopped at indicated time points by addition of 2X sample buffer and boiling for 5 min. The reaction mixture (∼150ng of Rab10) was loaded on 12% polyacrylamide gel. Samples were detected with anti-Rab10 and anti-pRab10 antibodies as indicated. Bottom, quantification of pRab10/ Rab10 signal. (B) As mentioned above, Purified Rab10 Q63L and Mst3 were incubated together and supplemented ± 2 mM ATP. Reactions were stopped, loaded on 12% SDS-polyacrylamide gel and analyzed as in A. (C) Purified Rab10-Q63L was mixed with purified Mst3 in reaction buffer as indicated and phosphorylated at 27°C for 2h. The reaction mixture (∼200-250ng of Rab10) was loaded onto a Phostag gel (10% gel with 40μM Phostag, 80μM MnCl_2_) and transferred onto nitrocellulose membrane after EDTA washes and probed with anti-Rab10 antibody. Two separate Phostag gels are shown. (D) Schematic of pRab10 phosphorylation, binding conditions and MST experiments. In +ATP condition, Rab10 QL and Mst3 are incubated for 2 h at 27°C. Pre-spun NHS-Red Labeled GTD (final 100nM) is then added to serially diluted pRab10 QL reaction mixture and incubated for 30 min in the dark at 25°C before MST is started (top). In -ATP condition, the reaction mixture of Rab10 QL and Mst3 are serially diluted, and NHS-red labeled GTD (final 100nM) is added immediately. The samples are incubated for 30min in the dark at 25°C before MST measurements are started. In all phosphorylation reactions, final concentrations of Rab10 and Mst3 were 20μM and 7μM respectively.

Because it is difficult to purify significant quantities of fully active and phosphorylated Rab10, we established a protocol that would enable us to monitor pRab10 binding to MyoVa with minimal protein handling. As outlined in Fig. 7D, Rab10 was incubated for 120 minutes with Mst3 kinase and ATP to achieve phosphorylation. NT-647 dye-labelled GTD was added for a subsequent 30 minute binding reaction, followed by transfer of the binding mix into capillaries for microscale thermophoresis analysis. Control reactions contained all the same components except ATP during binding; no signal was observed in reactions lacking Rab10 protein.

Phosphorylated Rab10 bound to purified MyoVa GTD with a K_D_ of 475 ± 166 nM under these conditions (Fig. 8A). This represents strong binding as most Rab effectors bind with low micromolar affinities. Non-phosphorylated Rab10 assayed in parallel showed no binding to GTD in reactions containing up to 18 μM GTD (Fig. 8B). As a positive control, Rab11A also bound to the MyoVa GTD with a K_D_ of ∼246 ± 216 nM (Fig. 8C). This affinity is similar to that previously reported for Rab11A binding to the closely related Myosin Vb GTD (Pylypenko *et al*, 2013). In summary, these in vitro binding assays confirm that pRab10 binds directly and with high affinity to the MyoVa GTD.

**Figure 8.**
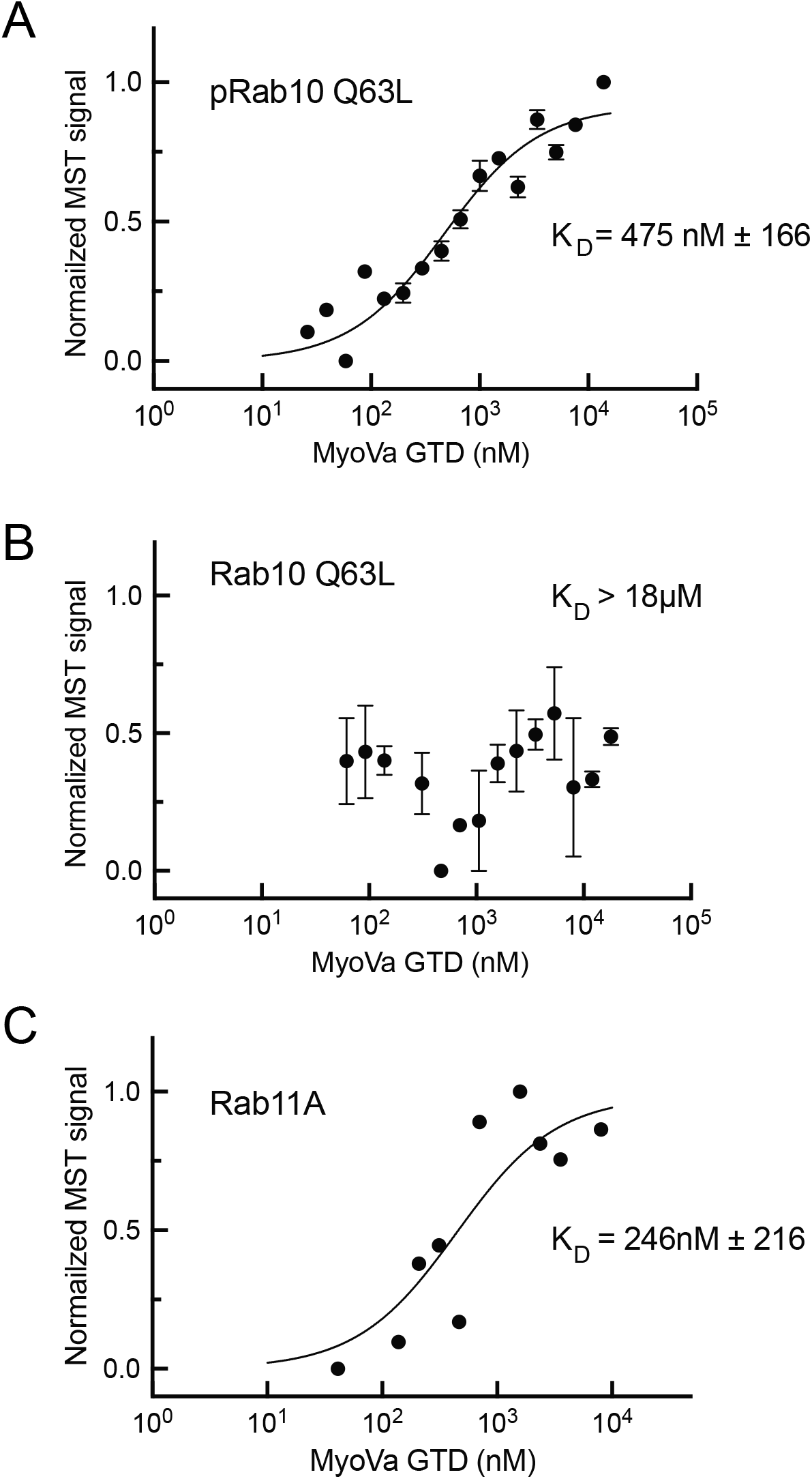
pRab10 binds the MyoVa globular tail domain. (A) Microscale thermophoresis (MST) of the interaction between labeled GTD with phosphorylated Rab10 Q63L (1-181). Purified Rab10 Q63L (1-181) was phosphorylated with Mst3 at 27°C for 2h and then serially diluted (26-13800 nM)). Immediately after, NHS-RED labeled GTD (final concentration 100nM) is added. (B) MST of Rab10 binding to unphosphorylated Rab10 Q63L. (C) MST of WT Rab11 binding to GTD measured in the absence of Mst3 kinase. Graphs in A and B show mean and SEM from two independent experimental measurements, each from a different set of protein preparations.

### Exogenous RILPL2 localizes to the mother centriole and blocks ciliogenesis

We showed previously that overexpression of RILPL1 was sufficient to block ciliogenesis in RPE and A549 cells (Dhekne *et al*, 2018). In addition, LRRK2 generation of pRab10 was sufficient to block ciliogenesis but this required that cells contain endogenous RILPL1 protein (Dhekne *et al*, 2018). We thus tested the effect of exogenous RILPL2 expression on primary cilia formation. hTERT-RPE cells were transfected with HA-tagged RILPL2 and 24 hours later, serum starved overnight to initiate ciliogenesis. As shown in Fig. 9 (A,B), even in the absence of pathogenic LRRK2, RILPL2 expression strongly inhibited cilia formation. In addition, clonal RILPL1 and RILPL2 knockout A549 cell lines were created using CRISPR. As we reported previously (Dhekne *et al*, 2018), endogenous RILPL1 depletion in these cells dramatically increased cilia formation, consistent with its role as a suppressor of ciliogenesis. In contrast, RILPL2 depletion did not influence ciliation in two independent knockout lines (Fig. 9C).

**Figure 9.**
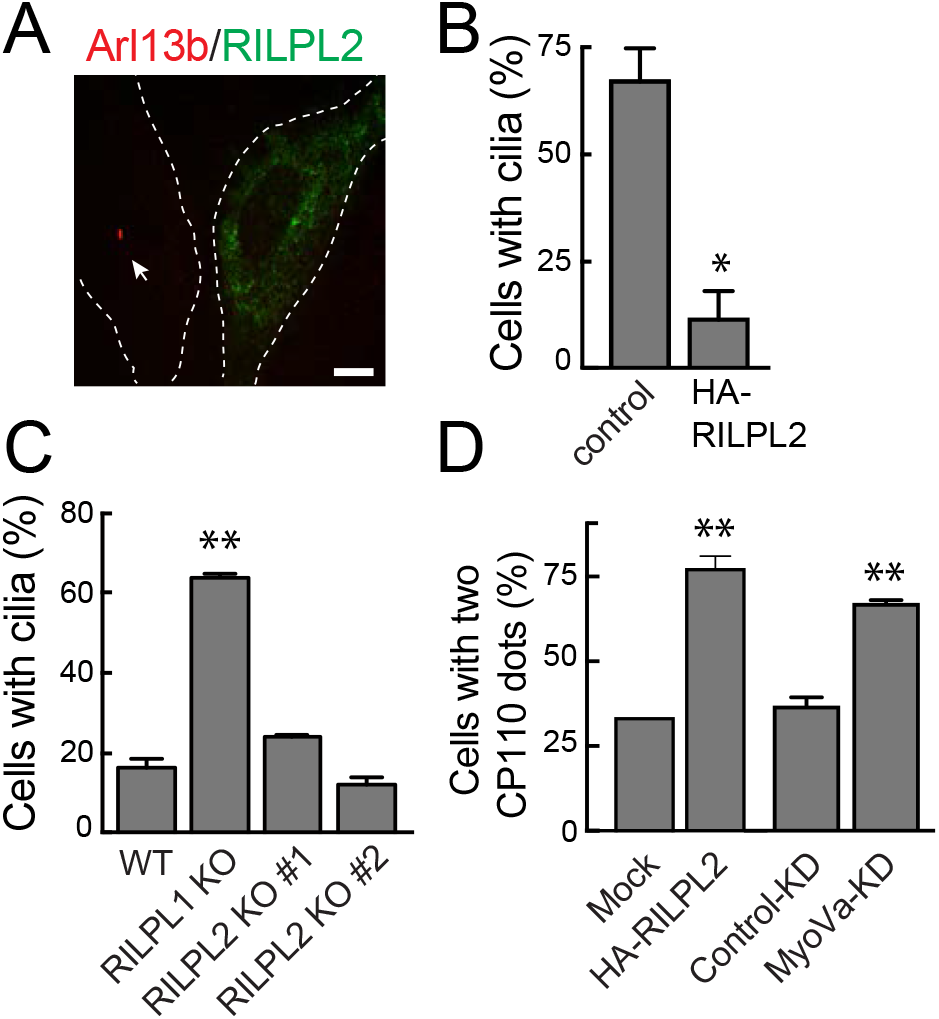
Exogenous RILPL2 blocks ciliogenesis CP110. (A) hTERT-RPE cells were transfected with HA-RILPL2 or mock transfected with lipofectamine and 24h later, serum starved overnight to initiate ciliogenesis. Cells were fixed with ice cold −20°C methanol for 2min and co-stained with rabbit anti-HA (green) or mouse anti-Arl13b (red). Arrow indicates the presence of an Arl13b^+^ cilium. Dotted lines indicate cell outlines. Bar, 10 μm. (B) Quantitation of cells with cilia ± HA-RILPL2. Error bars represent SEM from two experiments, each with >25 cells per condition. Significance was determined by the t-test; *, p = 0.0164. (C) WT, RILPL1, and RILPL2 knockout A549 cells were plated at 80% confluency. After 24h, cells were serum starved by transfer into 2% FBS-containing medium for 48h. Cells were fixed and stained with mouse anti-Arl13b. Quantitation of cells with cilia. Error bars represent SEM from two experiments with >100 cells per condition in each experiment. #1 and #2 indicate two different A549 RILPL2 knockout clones. (D) Quantitation of cells with CP110 dots indicating capped centrioles in: (left) hTERT-RPE cells ± HA-RILPL2 for 24 h; (right) hTERT-RPE cells ± MyoVa-siRNA for 72h. Cells were serum starved for 2h, fixed with −20°C methanol for 2 min and co-stained with anti-HA, anti-CP110 or anti-CEP164/CP110. Error bars represent SEM from two experiments with >25 cells per condition in each experiment. Significance was determined by the t-test; **, p = 0.0067 and 0.0045.

Ciliogenesis is initiated by Tau tubulin kinase 2 (TTBK2)-mediated phosphorylation events that drive the uncapping of CP110 from the mother centriole (Spektor *et al*, 2007; Goetz *et al*, 2012). To further explore the mechanism of RILPL2-mediated cilia blockade, we investigated the effect of RILPL2 expression on centriolar CP110 cap removal upon serum withdrawal. As shown in Figure 9D, fewer cells expressing RILPL2 lost CP110 from their mother centrioles, indicating that RILPL2 expression inhibits ciliogenesis at an early stage, upstream of TTBK2. CP110 also failed to be released from the mother centriole in MyoVa-depleted cells within 2 hours of serum starvation (Fig. 9D), a condition that leads to redistribution of RILPL2 to the mother centriole (Figure 2C,D).

Wu *et al* (2018) reported that MyoVa knockout cells showed no effect on CP110 release but had a striking defect in ciliary vesicle association with mother centriole. This difference might be explained by adaptation in knockout clones and/or a very short time of serum starvation (30 mins) in Wu *et al* (2018, Supp Fig. 4B) versus acute, siRNA-mediated MyoVa depletion and longer serum starvation (2 hours in serum free medium) in our experiments (Fig. 9D). Upon serum starvation, pre-ciliary vesicles fuse to form a ciliary vesicle in an EHD1 dependent manner, a step that is essential for CP110 uncapping and subsequent ciliogenesis (Lu *et al*, 2015). Consistent with a role in trafficking pre-ciliary vesicles in early ciliogenesis (Wu et al, 2018), MyoVa depletion in our experiments blocked CP110 uncapping. It is possible that tight binding to either exogenously expressed RILPL2 or pRab10 generated by LRRK2 sequesters MyoVa in a manner that cannot carry out its normal roles in cilia formation.

Altogether, the experiments presented here suggest that RILPL2 is usually removed from the centriolar region by MyoVa, unless cells express pathogenic LRRK2 and generate significant levels of peri-centriolar pRab10 protein. Under those conditions, or conditions of RILPL2 overexpression, RILPL2 and MyoVa accumulate at the centriole concomitant with a block in the formation of primary cilia. Moreover, the absence of MyoVa exon D in the brain is predicted to lead to the greatest change in MyoVa localization when pRab10 is present.

## Discussion

RILPL2 binds LRRK2-phosphorylated Rab10 with greater affinity than non-phosphorylated Rab10 (Steger *et al*, 2017), and although RILPL2 is known to bind MyoVa (Lisé *et al*, 2009) and contribute in some way to cargo enrichment in ciliogenesis (Schaub & Stearns, 2013), little is known about its precise cellular function. We have shown here that RILPL2 is diffusely localized in RPE cells, and a small portion moves to the mother centriole within ∼3 hours of serum starvation (Figs. 10A,B). The dispersed localization of RILPL2 is controlled by its MyoVa binding partner: depletion of MyoVa tripled the percent of cells displaying centriolar RILPL2 protein (Fig. 10C). This suggests that MyoVa functions at least in part to transport RILPL2 away from the mother centriole. In contrast, when expression of pathogenic R1441G LRRK2 increases pRab10 at the mother centriole, RILPL2 levels also increase there, and additional co-expression of low levels of MyoVa is sufficient to drive essentially all RILPL2 to the centriole (Fig. 10D). MyoVa is normally broadly distributed throughout the cytoplasm but it also concentrated at the mother centriole when pRab10 levels are high. In addition, peri-centriolar concentration of MyoVa and RILPL2 both require Rab10 protein.

**Figure 10.**
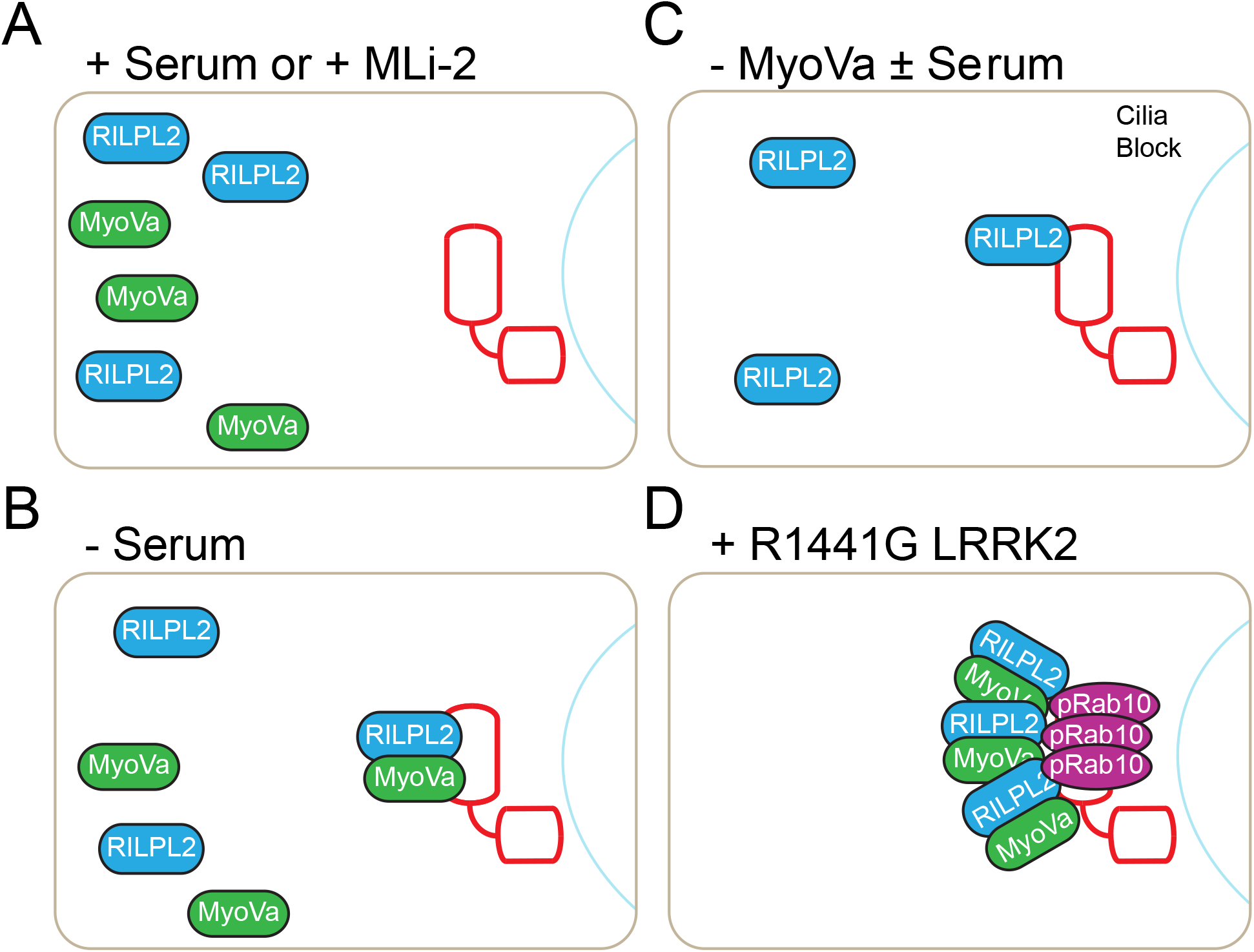
Summary of Protein Distributions. (A) Normal serum growth conditions or with LRRK2-specific MLi-2 inhibitor in cells expressing R1441G LRRK2; (B) Upon serum removal to trigger ciliogenesis; (C) In cells depleted of MyoVa using siRNA; (D) Upon expression of R1441G LRRK2 kinase.

Our studies revealed that pRab10 interacts directly and tightly with MyoVa’s globular tail domain, and this interaction is likely responsible for the relocalization of MyoVa to the centriole in R1441G LRRK2-expressing cells. This was unexpected, because pRab10 could have easily influenced MyoVa localization indirectly via RILPL2 binding. The globular tail domain of MyoVa is a versatile cargo adaptor with independent binding sites for RILPL2, Rab11A, Rab3A and melanophilin. Future experiments will determine the precise interface between pRab10 and MyoVa and whether or not pRab10 binding precludes interaction with additional binding partners. That MyoVa is a Rab11 effector suggests that it will function as part of the Rab11:Rab8 cascade that is an important part of the early steps of cilia formation (Knödler *et al*, 2010; Franco *et al*, 2014).

Our experiments indicate that MyoVa lacking exon D essentially binds only to phosphorylated Rab10 protein. Because expression of MyoVa lacking exons D and F is specific to endocrine, neuroendocrine and neuronal cells (Seperack et al, 1995), Rab10 phosphorylation becomes a primary mode of regulation for MyoVa in these cell types. Thus, the RH homology domain proteins RILPL1, RILPL2, JIP3 and JIP4 increase their binding to Rab10 upon LRRK2 phosphorylation, while MyoVa lacking exon D engages a new partner in the brain and endocrine cells. This may also have cell type-specific implications for MyoVa motor activity (Cao *et al*, 2019).

Despite its ability to bind RILPL2, endogenous Rab8 which is also phosphorylated by R1441G LRRK2 and located at the centriole (Purlyte et al., 2018; Ordonez *et al*, 2019) does not appear to be sufficient to drive RILPL2 relocalization in cells depleted of Rab10. This may be because MyoVa binds with preference to Rab10, and the MyoVa globular tail domain binds with strong preference to pRab10 and apparently not to pRab8 (Waschbüsch *et al*, 2020).

Knockout of RILPL2 does not alter the ability of cells to form primary cilia. This is unlike what we reported previously for RILPL1: loss of RILPL1 enhances cilia formation, as if it’s normal role is to repress this process (Dhekne *et al*, 2018). Nevertheless, like RILPL1, expression of HA-RILPL2 blocks ciliogenesis, perhaps by sequestering important components (such as MyoVa or Rab10) in a manner that interferes with their ability to function. Interestingly, video microscopy of protein recruitment to the mother centriole upon serum starvation shows that fluorescently tagged MyoVa arrives within 30 min (Wu et al 2018) whereas GFP-RILPL2 required more than two hours. This points to a possible lead role of centriolar Rab10 in recruiting MyoVa protein. As for RILPL2, its recruitment to the centriolar region could reflect binding to peri-centriolar vesicle-associated Rab10, MyoVa, or both.

PhosphoRab10 represents a potent interaction partner for RILPL1, RILPL2 and MyoVa globular tail domain. Interestingly, all of these proteins are implicated in some way in the normal process of ciliogenesis and all become clustered around the mother centriole in cells expressing pathogenic LRRK2 kinase that are blocked in ciliogenesis. Are any of these components more important than another in regulating cilia formation? In our previous work, we showed that RILPL1 and Rab10 are essential components of the LRRK2-mediated, ciliogenesis blockade (Dhekne *et al*, 2018). Here we show that loss of RILPL2 has no consequence for cilia formation, while RILPL1 depletion significantly enhances cilia formation. According to mapofthecell.biochem.mpg.de, RILPL1 is present at 72,522 copies per HeLa cell and 98% cytosolic, compared with 45,719 copies of MyoVa, which fractionates most significantly with endosomal markers (Itzhak *et al*, 2016). However, RILPL2 was below detection in this tour de force mass spectrometry experiment (Itzhak *et al*, 2016). Similarly, the ratio of RILPL2 to RILPL1 is quite variable across tissues (pax-db.org) and cell lines and thus differences in the contributions of RILPL1 and RILPL2 to ciliogenesis blockade may reflect their relative abundance in different cell types. Unlike RILPL2, RILPL1 does not bind MyoVa. As described herein, any role for RILPL2 in blocking ciliogenesis likely requires MyoVa participation since it so significantly altered RILPL2’s cellular distribution.

Although it is not yet known how RILPL2 becomes localized to the mother centriole upon serum starvation, it’s expression at higher than endogenous levels is coupled with a block in the release of the CP110 cap from the mother centriole upon serum starvation. Release of the CP110 cap by the action of Tau tubulin kinase 2 and EHD1/3 dependent ciliary vesicle formation are essential steps in the process of ciliogenesis (Spektor *et al*, 2007; Goetz *et al*, 2012; Lu *et al*, 2015). MyoVa depletion also blocks release of CP110 in our experiments, and this may be explained in part, by the increase in peri-centriolar RILPL2 seen in cells depleted of MyoVa. Indeed, MyoVa strongly regulated RILPL2’s localization, consistent with their tight molecular association and cooperative roles. In the presence of pathogenic LRRK2 kinase, MyoVa and pRab10 drive a major relocalization of RILPL2 to the mother centriole, and it will be important to explore further how this contributes to a pathogenic LRRK2-triggered ciliogenesis blockade.

## Materials and Methods

### Reagents

MLi-2 LRRK2 inhibitor was obtained from Tocris Biosciences (Cat. No. 5756).

### General cloning and plasmids

DNA constructs were amplified in *E. coli* DH5α or stbl3 and purified using mini prep columns (Econospin). DNA sequence verification of all plasmids was performed by Sequetech (http://www.sequetech.com). Myc-LRRK2 was obtained from Addgene (#25361) and the R1441G mutation introduced as described (Purlyte *et al*, 2017). RILPL2-GFP was cloned as described (Steger *et al*, 2017). N-terminally tagged GFP-RILPL2 was cloned into pcDNA5D using Gibson assembly while HA-RILPL2 was cloned into BamH1 and Not1 sites of pcDNA5D by restriction cloning. Mouse MyoVa-mCherry plasmid was a gift of Prof. Jim Spudich (Stanford University). MyoVa-GTD-mCherry (1421-1880) and exon D (1320-1345) deletion mutants were made by site directed mutagenesis. For bacterial expression, MyoVa GTD, Rab10-Q63L (1-181) and Mst3 were kind gifts from Amir Khan (Trinity College, Dublin, Ireland). mKO2-PACT was generated by amplifying the PACT domain of Pericentrin from cDNA of RPE cells as described (Sobu *et al*, 2020).

### Cell culture, transfections and ciliogenesis

HEK293T, A549, and hTERT-RPE cells were obtained from ATCC cultured in high glucose DMEM with 10% fetal bovine serum. HEK293T cells were transfected with Polyethylenimine HCl MAX 4000 (PEI) (Polysciences, Inc.) as described (Reed *et al*, 2006). RPE cells were transfected with Fugene HD (Promega) while A549 cells were transfected using Lipofectamine 3000 according to the manufacturer. RPE cells stably expressing mKO2-PACT were generated by lentivirus as described at (Sobu *et al*, 2020). Cells were checked routinely for Mycoplasma using either MycoALert Mycoplasma Detection Kit (Lonza LT07-318) or PCR. Cells were grown to confluence in DMEM with serum. hTERT-RPE cells were ciliated by overnight serum starvation in DMEM medium alone. For A549 cells, cells were plated at 80% confluency and 24h later, subjected to 2% serum for 48h. To identify primary cilia, cells were stained for a cilia marker, Arl13b. For monitoring CP110 release, RPE cells were serum starved for 1h. Cultures of primary rat astrocytes were obtained by antibody panning from rat pups as described (Foo *et al*, 2011; Dhekne *et al*, 2018). In brief, six to ten postnatal Sprague-Dawley rat cortices were mechanically and enzymatically dissociated to produce single cells. They were passed over successive, antibody-coated negative panning plates to rid the suspension of microglia, endothelial cells, and oligodendrocyte precursor cells before selecting for astrocytes with an anti-ITGB5-coated plate. Astrocytes that attached to the anti-ITGB5-coated plate were trypsinized and plated onto poly-D lysine coated ACLAR coverslips. Cells were incubated at 37°C with 10% CO_2_ and 90% O_2_ for one to three weeks.

### Lentivirus production

Lentivirus-based shRNA was used for gene knockdown of human Rab10. The lentiviral vector was co-transfected with packaging vectors psPAX2, pMD2 VSV-G in HEK293T cells using PEI. pLKO-puro-scramble was used as control. After 48 h, culture supernatants were collected and virus concentrated 10x overnight with Lenti-X concentrator (Clontech) according to the manufacturer. Cells were transduced with virus using polybrene (2 μg/ml). To make stable cells, infected cells were selected with puromycin (0.2–1 μg/ml) 48 h after infection. Expression of the target protein was verified by qPCR and/or immunoblotting.

### Gene Knockdown

For silencing of the Rab10 gene, lentivirus pLKO.1-Rab10 shRNA was used to infect HEK293T cells as described (Dhekne *et al*, 2018). For silencing MyoVa, siRNA against MyoVa was obtained from Invitrogen (s9207, 5’ GUAUAGUCCUAGUAGCUA 3’) as this was shown to work well by Wu *et al* (2018), and transfected at 30 nM final concentration using Lipofectamine 3000 according to the manufacturer. After 48h, cells were plated onto coverslips for ciliogenesis or processed to determine knock-down efficiency by qPCR. After 72h, cells were lysed for western blot analysis of the MyoVa knock-down.

### Lysing and Immunoblotting

Cells were lysed 24h after transfection in ice-cold lysis buffer containing 50 mM Tris/HCl, pH 7.5, 150 mM NaCl, 1% (v/v) Nonidet P-40 (NP-40), 5 mM MgCl_2_, 5 mM ATP, 1 mM EGTA, 1 mM sodium orthovanadate, 50 mM NaF, 10 mM 2-glycerophosphate, 5 mM sodium pyrophosphate, 0.1 μg/ml mycrocystin-LR (Enzo Life Sciences), 1 mM DTT and EDTA-free protease inhibitor cocktail (Sigma). Lysates were centrifuged at 13800 *x g* for 15 min at 4°C and supernatants quantified by Bradford assay (Thermo Scientific) and 60 μg and 400 μg of proteins used for immunoblotting and coimmunoprecipitation assay, respectively. Antibodies were diluted in a blocking buffer containing 5% milk in 0.05% Tween in Tris buffered saline (TBS) and incubated overnight on the blots. Phos-tag (Fujifilm Wako) gels were made according to the manufacturer using 40 μM Phostag, 80 μM MnCl_2_ in a 10% acrylamide-bis-acrylamide gel. The phostag gel was washed for 15 min in 10 mM EDTA/Water, followed by 5 min wash in 1 mM EDTA/water and finally, distilled water. Phosphorylated proteins were transferred to nitrocellulose for 20 min using a Biorad Transblot system. Lysis for qPCR was performed by solubilizing cells in 500 μl Trizol. RNA was extracted according to the manufacturer. cDNA was synthesized using Expand Multiscribe cDNA synthesis kit (4368814) by Applied Biosystems. 1:10 diluted cDNA was then subjected to qPCR using Powerup Sybr green master mix (Thermo) in an Applied Biosystem ViiA 7 qPCR machine. qPCR primers were from Origene (MyoVa Fw - CTCACACGAACTCCTGCAAA, MyoVa Rv - AGGGGTAGTGGCATTGAGTG). The mRNA expression analysis was performed using the delta delta Ct method.

### Co-Immunoprecipitation

HEK293T cells expressing RILPL2-GFP, MyoVa-mCherry or myc-LRRK2 R1441G were harvested 24–48 h post-transfection and lysed. Equal amounts of extract protein were incubated with GFP-binding protein-conjugated agarose (Chromotek) or anti-RFP antibody (Anti-RFP RTU - Rockland Biosciences) conjugated protein G beads (Thermo) for 1 h at 4°C. Immobilized proteins were washed 4 times with 1 ml lysis buffer, eluted with 2X SDS loading buffer, and subjected to BioRad Mini-PROTEIN TGX 4-20% gradient gels. After transfer to nitrocellulose membrane and antibody incubation, blots were visualized using the LI-COR Odyssey Imaging System.

### Generation of knockout A549 cells

Rab10 knock-out (Ito *et al*, 2016) cell lines have been described (Steger *et al*, 2017). The A549 RILPL1 knockout cell line has been described (Dhekne *et al*, 2018) and the RILPL2 knockout cell lines were made similarly using these guide pairs: sense A: GCCTGTTGGGCCGCGAGCTTA; antisense A: GATGTCATACACGTCCTCGG and sense B: GTATGACATCTCCTACCTGTT, antisense B: GCCTCGGCGGTCAGCTGGAAG.

### Immunofluorescence staining and light microscopy

Cells were plated on collagen coated coverslips transfected with indicated plasmids. Cells were fixed with 3% paraformaldehyde for 20 min, permeabilized for 3 min in 0.1% Triton X 100 (or 0.1% saponin for anti-pRab10 antibody staining) and blocked with 1% BSA in PBS. To stain centrioles with anti-CP110, anti-CEP164, or anti-EHD1, cells were fixed with ice-cold 100% methanol for 5 min and gently washed with PBS followed by blocking with 1% BSA in PBS.

Antibodies were diluted as follows: mouse anti-Arl13b (1:2000, Neuromab), rabbit anti-Arl13b (1:1000, Proteintech); mouse anti-GFP (1:2000, Neuromab); rabbit anti-RFP (1:1000, Rockland); mouse anti-Myc (1:2, 9E10 Hybridoma culture supernatant, 1:1000 Biolegend); rabbit anti pRab10 (1:1000, Abcam); mouse anti-CEP164 (1:1000, Santa Cruz); rabbit anti-CP110 (1:2000, Proteintech); rabbit anti-EHD1 (1:1000, Abcam), rabbit anti-MyoVa (1:1000, Cell Signaling Technology), rabbit anti-LRRK2 (Clone UDD3, Abcam), rabbit anti-RILPL2 (1:500, Novus) and mouse/rabbit anti-HA (1:1000, Sigma). Anti-sera raised against RILPL2 in rabbit is described in Steger *et al* (2017) was used for staining immuno-panned primary rat astrocytes. Highly cross absorbed H+L secondary antibodies (Life Technologies) conjugated to Alexa 488, Alexa 568 or Alexa 647 were used at 1:2000 or 1:4000. Nuclei were stained using 0.1 μg/ml DAPI (Sigma).

All images were obtained using a spinning disk confocal microscope (Yokogawa) with an electron multiplying charge coupled device (EMCCD) camera (Andor, UK) and a 100X 1.4NA oil immersion objective. Images were analyzed using Fiji (https://fiji.sc/) and as indicated, are presented as maximum intensity projections. Colocalization was quantified using the plugin JaCoP. Pearson’s and Mander’s correlation coefficients were used to quantify colocalization between fluorophores.

For time lapse imaging, 3×10^4^ hTERT-RPE cells expressing mKO2-PACT were seeded into 8 well glass bottom dishes (Nunc^™^ Lab-Tek^™^ II Chambered Coverglass, Thermo Fisher Scientific) and cultured overnight. Cells were transfected with GFP-RILPL2, then, after 24h cells were serum starved by replacing media with Leibovitz’s L-15 Medium, no phenol red (Thermo Fisher Scientific) without serum. Starting 15 min after starvation initiation, images were captured every 6 min using a spinning disk confocal with 63X glycerol immersion objective at 37°C.

### Protein expression and purification

His_6_ Rab10-Q63L (1-181), His_6_ Mst3, His_6_ Myosin Va GTD (1464-1855), and His_6_ Rab11a were purified in *E. coli* BL21 (DE3 pLys) as described by Waschbüsch *et al* (2020) and Berndsen *et al* (2019). In brief, bacterial cells were grown at 37°C in Lucia Broth medium and induced at A_600_ nm = 0.6-0.7 by the addition of 0.1 mM isopropyl-1-thio-□-D-galactopyranoside (Gold Biotechnology) and harvested after 18h at 18°C. The cell pellets were resuspended in ice cold lysis buffer (50 mM Tris pH 8.0, 10% (v/v) glycerol, 250 mM NaCl, 10 mM Imidazole, 5 mM MgCl_2_, 0.5mM DTT, 20 □M GTP, and EDTA-free protease inhibitor cocktail (Roche)), lysed by one passage through an Emulsiflex-C5 apparatus (Avestin) at 10,000 lbs/in^2^, and centrifuged at 13,000 rpm for 25 min in a FiberLite F15 rotor (ThermoFisher). Clarified lysate was incubated with cOmplete Ni-NTA resin (Roche) for 2h at 4°C. Resin was washed three times with 5 column volumes lysis buffer and eluted in 500 mM imidazole-containing lysis buffer. The eluate was buffer exchanged and further purified by gel filtration on Superdex-75 (GE Healthcare) with 50 mM Hepes pH 7.5, 5% (v/v) glycerol, 150mM NaCl, 5mM MgCl_2_, 0.1 mM tris(2-carboxyethyl)phosphine (TCEP), and 20 □M GTP.

### In vitro phosphorylation and microscale thermophoresis (MST)

His_6_ Rab10 Q63L 1-181 was incubated with His_6_-Mst3 at a molar ratio of 3:1 (substrate: kinase). The reaction buffer was 50 mM Hepes pH 7.5, 5% (v/v) glycerol, 150 mM NaCl, 5mM MgCl_2_, 0.1 mM TCEP, 20 mM GTP, 6 □M BSA, 0.01% Tween-20, and 2 mM ATP (no ATP for negative control). The reaction mixture was incubated at 27°C for 30 min – 4.5 h in a water bath. Phosphorylation completion was assessed by western blot and PhosTag gel electrophoresis and found to saturate at 2 h. Immediately following phosphorylation, samples were transferred to ice prior to binding determination.

Protein-protein interactions were monitored by MST using a Monolith NT.115 instrument (Nanotemper Technologies). His_6_-MyoVa GTD was labeled using RED-NHS 2^nd^ Generation (Amine Reactive) Protein Labeling Kit (Nanotemper Technologies). For all experiments, the unlabeled protein partner was titrated against a fixed concentration of the fluorescently labeled GTD (100 nM); 16 serially diluted titrations of the unlabeled protein partner were prepared to generate one complete binding isotherm. Binding was carried out in reaction buffer in 0.5mL Protein Lobind tubes (Eppendorf) and allowed to incubate in the dark for 30 min before loading into NT.115 premium treated capillaries (Nanotemper Technologies). A red LED at 10% excitation power (red filter, excitation 605–645 nm, emission 680–685 nm) and IR-laser power at 20% was used for 30 sec followed by 1 sec of cooling. Data analysis was performed with NTAffinityAnalysis software (NanoTemper Technologies) in which the binding isotherms were derived from the raw fluorescence data and then fitted with both NanoTemper software and GraphPad Prism to determine the *K_D_* using a non-linear regression method. The binding affinities determined by the two methods were similar. Shown are averaged curves of pRab10 and Rab10 from single readings from two different protein preparations. Rab11 positive control curve is from a single reading from a single protein preparation due to COVID-19 shutdown.

### Statistics

Graphs were made using Graphpad Prism 6 software. Error bars indicate SEM. Unless otherwise specified, a Student’s T-test was used to test significance. Two tailed p-values < 0.05 were considered statistically significant.

## Acknowledgements

This research was funded by a grant to SRP from the US NIH (DK37332), and grants from the Michael J Fox Foundation for Parkinson’s research. We are grateful to Dr. Dario Alessi for critical input and enthusiastic support.

## Author Contributions

IY, HSD, EGV, YS carried out the experiments presented herein and wrote and edited the manuscript; FD created knockout cell lines and SRP oversaw the research, wrote and edited the paper and obtained funding to support the study.

